# Th17/regulatory T cells balance is predictive of *Coccidioides* infection outcome in pediatric patients

**DOI:** 10.1101/347898

**Authors:** Dan Davini, Fouzia Naeem, Aron Phong, Mufadhal Al-Kuhlani, Kristen M. Valentine, James McCarty, David M. Ojcius, David M. Gravano, Katrina K. Hoyer

## Abstract

**Background:** Protective immunity against the fungal pathogen Coccidioides requires specific T helper responses. Mouse vaccine and infection studies have defined CD4^+^ T helper (Th)1 and Th17 cells in the resolution of infection and in effective protection. Patients with persistent *Coccidioides* infection demonstrate reduced cellular responses.

**Methods:** Peripheral blood and serum were collected from 30 pediatric *Coccidioides*-infected patients and 20 healthy controls in the California San Joaquin Valley. Samples were evaluated by flow cytometry for innate and adaptive immune populations and cytokines to define the early immune response and identify clinically useful biomarkers for predicting disease outcome. Clinical and flow data were evaluated according to disease outcome (resolved or persistent) using principal component analysis, high-dimensional flow cytometry analysis tools, chi-square automatic interaction detection, and individual cell population comparisons.

**Results:** Patients with persistent infection had lower Th17 and higher Treg frequencies, but similar Th1 responses, relative to patients that resolved disease. Treg frequency, eosinophil numbers and neutrophil numbers together distinguish patients that resolve infection from those that develop persistent infection.

**Conclusions:** The inability to resolve *Coccidioides* infection may be a result of elevated Treg frequency and functional capacity, and Treg frequency may predict patient disease outcome at diagnosis. In our study, Th1 responses were similar in persistent and resolved infection, in contrast to prior human studies. Instead, our data suggest that Th17 cells provide an effective protection during *Coccidioides* infection, and that elevated Treg frequency inhibits protective immunity.

## INTRODUCTION

Coccidioidomycosis, also known as Valley fever, is caused by a fungal pathogen endemic to the San Joaquin Valley in California, Arizona, northern Mexico and arid areas of South America [1]. Population expansion, travel, improved detection and/or climate changes appear to be contributing to an expanding endemic region. 40% of those infected develop symptomatic pneumonia that typically resolves without anti-fungal therapy, although disease can become chronic and disseminate outside the lungs [1]. Persistent coccidioidomycosis is thought to be due to poor or ineffective cellular immunity.

Cellular immune responses are critical for effective immunity to *Coccidioides* infection [2, 3]. Early mouse studies indicate a need for T helper 1 (Th1) responses in protective immunity [4]. More recent studies highlight a role for Th17 cells and IL-17 cytokine responses [5–9]. Effective Th1 responses have been linked to resolution of human *Coccidioides* infection [10, 11]; however, Th17 effectors are critical for clearance of many other fungal pathogens.

Tregs function to control immune responses during infection and disease [12]. Th17 and Treg differentiation are inversely regulated, with IL-6 promoting TGFβ-induction of Th17 cells and inhibiting TGFβ-induction of Tregs [13–15]. An appropriate balance in T effector and regulatory cells allows effective immune responses while limiting tissue damage. Very little is known about the role of Tregs in the regulation of the immune response during *Coccidioides* infection. In mice infected with *Coccidioides posadasii*, Treg expansion correlated with reduced survival [16]. In other fungal infections, Treg expansion correlates with impaired T cell immunity and persistent disease [17–19]. IL-10, an anti-inflammatory cytokine produced by several adaptive immune cells including Tregs, promotes survival of *Coccidioides* and is associated with less-effective immunity [20, 21]. Tregs secrete IL-10 as one cellular immune suppression mechanism.

## METHODS

### Study enrollment and design

Patients were enrolled and samples collected at Valley Children’s Healthcare (VCH), a 355 bed Children’s Hospital serving as tertiary referral center for 10 counties in central California. Thirty children aged 2-18 years with coccidioidomycosis diagnosed by positive coccidioidal serology or cultures demonstrating *Coccidioides species* were included. Of these, 15 were enrolled as inpatients and 15 as outpatients at the time of diagnosis. Twenty healthy siblings of hospital patients, with negative coccidioidal serology were enrolled as controls. Children known to be pregnant, immunocompromised and/or on immunosuppressive medications, with severe underlying illness, cystic fibrosis, or inflammatory diseases were excluded. The study was approved by the VCH institutional review board. Written informed consent was obtained from legal guardians and participants >7 years of age. Baseline demographic, clinical, laboratory, radiographic, antifungal treatment and outcome data were collected. Outcomes were defined as resolved or persistent infection at the time of study conclusion. Resolution of symptoms, abnormal radiographic and clinical laboratory findings defined disease resolution. Persistent disease was defined as still on therapy without complete resolution of symptoms and/or persistent abnormal radiographic or laboratory findings.

Patients were characterized by 42 clinical parameters (Supplemental Table 1), 51 immune cell population parameters (Supplemental Table 2), and 26 serum proteins. Frequency and total number per milliliter of innate and adaptive immune populations were evaluated in the peripheral blood at the time of enrollment. We compared percentages, total number and activation state of immune cells in patients based on disease outcome (healthy control, resolved, persistent) and hospital status at time of blood draw (inpatient, outpatient).

### Serum and peripheral blood collection

1ml whole blood collected in sodium heparin tubes was stored at room temperature, and processed within 24-hours for flow cytometry. 1ml collected in red-cap plug tubes was centrifuged at 3500rpm for 15min then transferred into new tubes for −20°C storage.

### PBMC flow cytometric analysis

Antibody staining was performed in PBS/2%FBS. 100μL of whole blood was incubated with 5μL of Human TruStain FcX (Biolegend) for 5min prior to staining. Three flow cytometry panels were designed to profile PBMCs (Supplemental methods). A single blood sample was processed three times such that 50μL of antibody for each panel was added and incubated for 20min. RBCs were lysed in 1-step Fix/Lyse Solution (eBioscience) for 25min and resuspended with 300μL PBS/2%FBS and 100μL of 123count-eBeads (eBioscience) for flow cytometry analysis. Stained PBMC were acquired on an LSRII (BD), and analyzed using FCS Express (DeNovo Software) FACS Diva (BD).

To ensure consistent cytometer calibration and data collection, voltage and compensation parameters were standardized using SPHERO Ultra Rainbow Calibration Particle Kits (Spherotech, Inc.). Consistent MFI for each fluorophore was maintained by small adjustments to voltage parameters if baseline MFI changes were >10% between experiments. Fluorophore compensation was maintained with antibody single stains mixed from 1μL of each antibody into UltraComp eBeads (Bioscience), washed with PBS/2%FBS, incubated for 15min, and resuspended in 500μL PBS/2%FBS.

### High-dimensional flow cytometry analysis

FCS files were loaded into Cytobank (www.cytobank.org), and pre-gated to remove counting beads, debris, doublets, and dead cells. Remaining cells were gated on CD4+CD45+, grouped on outcome, and analyzed using CITRUS and viSNE. File internal compensation was utilized, clusters characterized on abundance, event sampling equalized per file, minimum cluster size set to 0.5%, and Significance Analysis of Microarrays (SAM) correlative model utilized. CITRUS analysis was repeated three independent times to confirm stratifying signatures and statistically significant populations (discovery rate <1%). viSNE map was generated from 373,781 total events (12,889/patient) using 2500 iterations, a perplexity of 90, and a Theta of 0.3.

### Cytokine assay

Thawed serum was centrifuged at 1000×*g* for 5min to remove particulates. Serum cytokine concentrations were determined using LEGENDplex Human 13-plex kits, ‘Th Cytokine Panel’ and ‘Cytokine Panel 2’ bead-based immunoassay kits (Biolegend) in duplicate according to the manufacturer’s instructions. Samples were diluted 2-fold with kit Assay Buffer prior to assay initiation. Samples were analyzed by flow cytometery and processed using LEGENDplex Data Analysis Software. Standard curves were generated using a 5-parameter curve-fitting model and cytokine levels were calculated as the average of the duplicate measurements.

### Data analysis and statistics

Leukocyte subset total numbers were calculated using counting beads in combination with each leukocyte percentage determined by flow cytometry. Statistical analyses were performed using Prism software v6.0 (GraphPad Software). One-way ANOVA with Bonferroni correction was used to compare multiple groups with a 95% confidence interval. The Fisher’s exact test was used to compare the distribution of groups and percentages. PCA and CHAID analyses were performed using XLSTAT.

## RESULTS

### Patient characteristics

Thirty pediatric patients with a diagnosis of coccidioidomycosis (15 inpatients and 15 outpatients) and 20 healthy controls were enrolled. One inpatient died due to disease severity and was excluded from analysis. Of the thirty patients with coccidioidomycosis, 25 had pulmonary involvement and five had disseminated disease including the deceased patient. Resolved disease was observed in 13 (44.8%) patients and persistent disease in 16 (55.2%), including 12 with pulmonary and 4 with disseminated disease.

Age and ethnicity, as well as the incidence of chest pain, night sweats, weight loss and fatigue symptoms were similar between the groups, and did not correlate with disease outcome (Table 1 and Table 2). More females than male patients were enrolled in the study, but this is unlikely to be due to an increased incidence in female children [22, 23]. Regardless of disease severity or outcome, most patients experienced fever. As observed previously, erythema nodosum mildly correlated with a better disease prognosis (p=0.0722). Evaluating complete blood count (CBC) measurements, inpatients had higher neutrophil, eosinophil, WBC and platelet numbers, and lower lymphocyte numbers compared to outpatients (Supplemental Figure 1). Eosinophil, platelet and creatinine (Cr) levels were elevated and lymphocyte counts reduced in patients with persistent disease compared to patients that resolved infection. Although some CBC parameters were significantly different between the groups, there was considerable overlap in values and these small variations would be difficult to use in predicting disease outcome. Serum IgM, IgG and antibody titers from *Coccidioides* enzyme immunoassay and complement fixation (CF) assays did not correlate with disease outcome, and were not different between inpatient and outpatients (Supplemental Table 3).

**Table 1.** Demographics of patients based on outcome status.

|  | Healthy Controls<br>(n=20) | Resolved<br>(n=13) | Persistent<br>(n=16) |
| --- | --- | --- | --- |
| Age (median, range), yrs | 11, 3-16 | 12, 6-18 | 13, 2-17 |
| Gender, n (%) |  |  |  |
| Male | 11 (54%) | 3 (23.1%) | 5 (31.3%) |
| Female | 9 (45%) | 10 (76.9%) | 11 (68.8%) |
| Ethnicity, n (%) |  |  |  |
| Hispanics | 14 (70%) | 11 (84.6%) | 14 (87.5%) |
| Non Hispanics | 6 (30%) | 2 (15.4%) | 2 (12.5%) |
| Hospitalization status, n (%) |  |  |  |
| Inpatients | 0 | 2 (15.4%) | 12 (75.0%) |
| Outpatients | 20 (100%) | 11 (84.6%) | 4 (25.0%) |

**Table 2.**
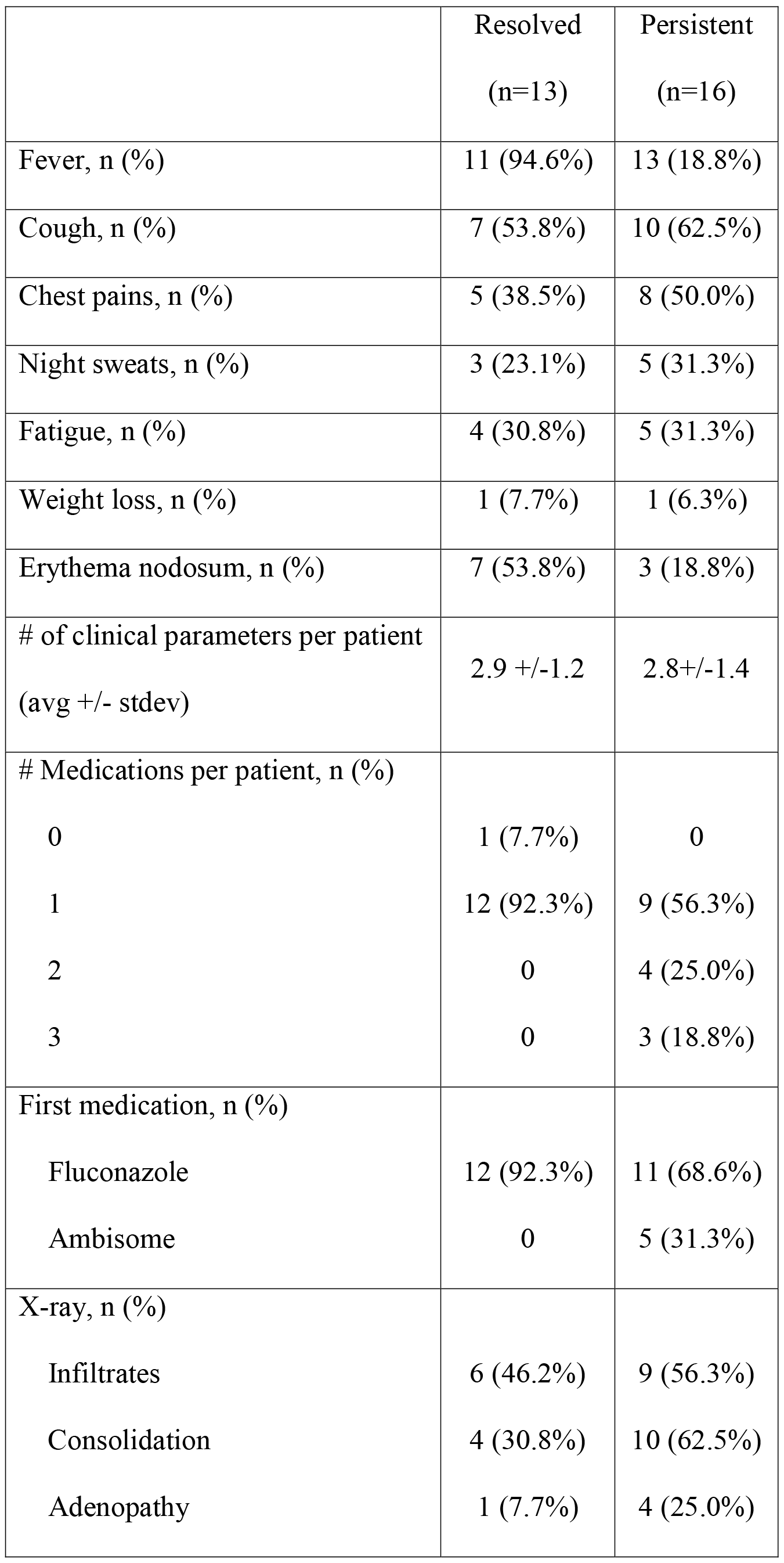
Clinical features of pediatric *Coccidioides*-infected patients.

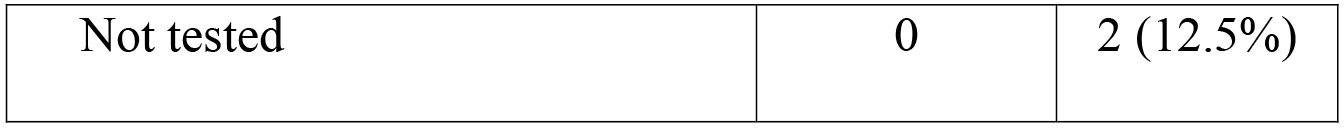

Analysis of immune parameters based on hospital status revealed several differences between inpatients and outpatients in both innate and adaptive responses. Plasmacytoid dendritic cell frequency was reduced in inpatients relative to healthy controls and outpatients (Figure 1A). CD4^+^, CD8^+^ and B cell frequency was reduced in inpatients, and mild elevation of Th1 was observed (Figure 1B, C). These differences likely reflect the more severe illness, elevated inflammatory responses and ineffective adaptive and Th cell immunity of the inpatients. A larger proportion of inpatients versus outpatients was unable to resolve *Coccidioides* infection, and included the four patients with disseminated disease, suggesting the more severe disease at diagnosis, the more likely that disease will persist.

**Figure 1.**
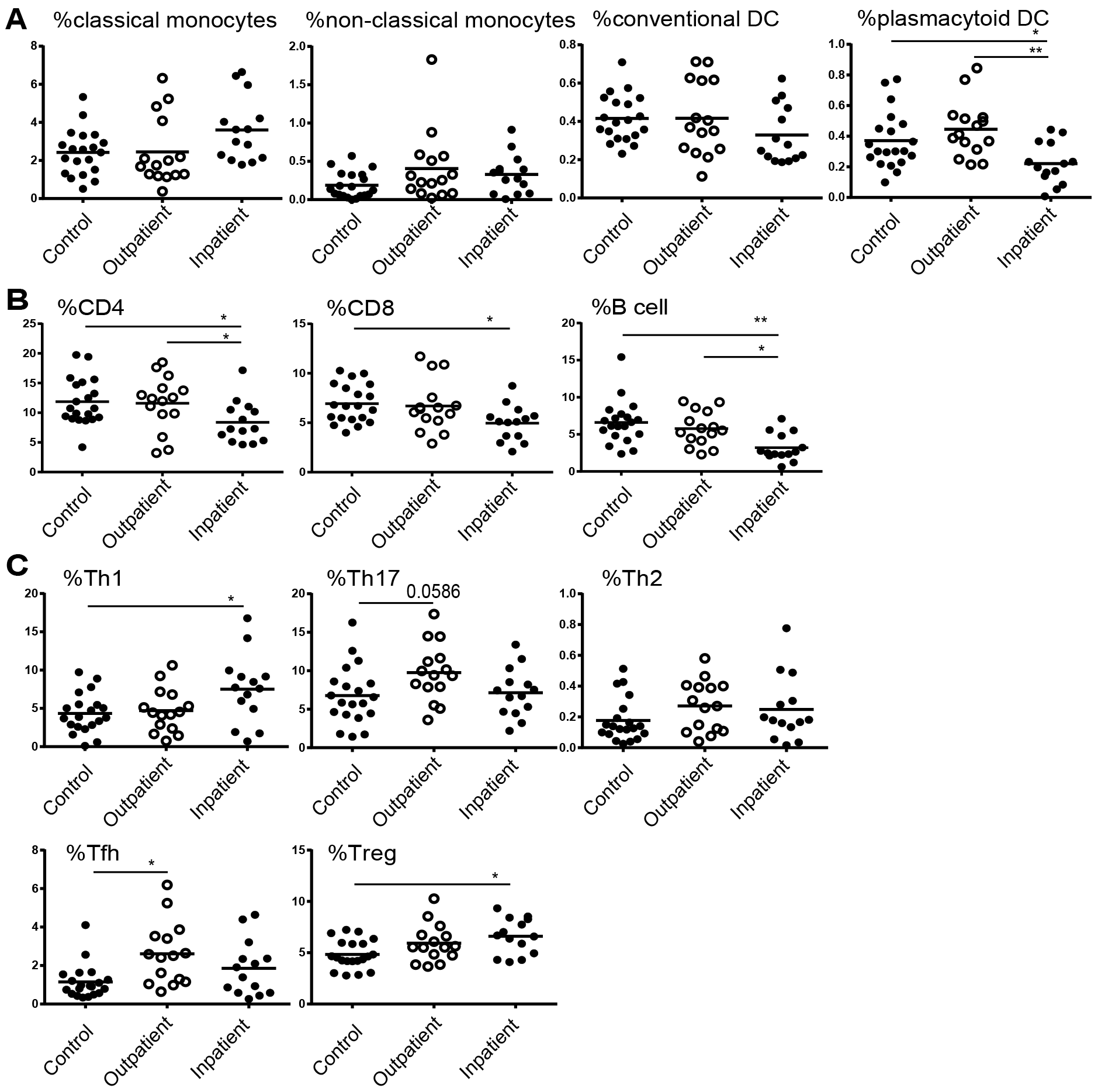
Immune parameters based on patient hospital status. PBMCs from healthy and *Coccidioides*-infected patients were assessed by flow cytometry. A) Comparisons of monocytes and dendritic cell frequencies. Frequency comparisons of adaptive immune populations CD4^+^, CD8^+^ and B cells (B), and of CD4^+^ T helper subsets as a percentage of CD4^+^ T cells (C). Each dot represents an individual patient, and lines indicate the mean. ANOVA with Bonferroni correction was used to compare multiple groups with a 95% confidence interval; * p<0.05, ** p<0.005, *** p<0.0005.

### Specific adaptive immune responses distinguish disease outcome

No differences were observed in the frequency or total number of innate immune populations based on disease outcome (data not shown). Patients with persistent disease tended to have lower adaptive immune cell frequency, in particular a significant reduction in B cell frequency relative to healthy controls and resolved patients (Figure 2A). Persistent patients also had fewer Tfh cells, important for effective and diverse antibody responses. Current paradigm suggests that a strong Th1 response is required for controlling *Coccidioides* infection. Patients with IFNγR or Stat1 mutations that reduce Th1 responses have more severe, disseminated infection [24]. Th1 frequencies were similar between patients with resolved and persistent disease (Figure 2B), while patients with persistent disease had lower Th17 and higher Treg frequencies than patients that resolved disease.

**Figure 2.**
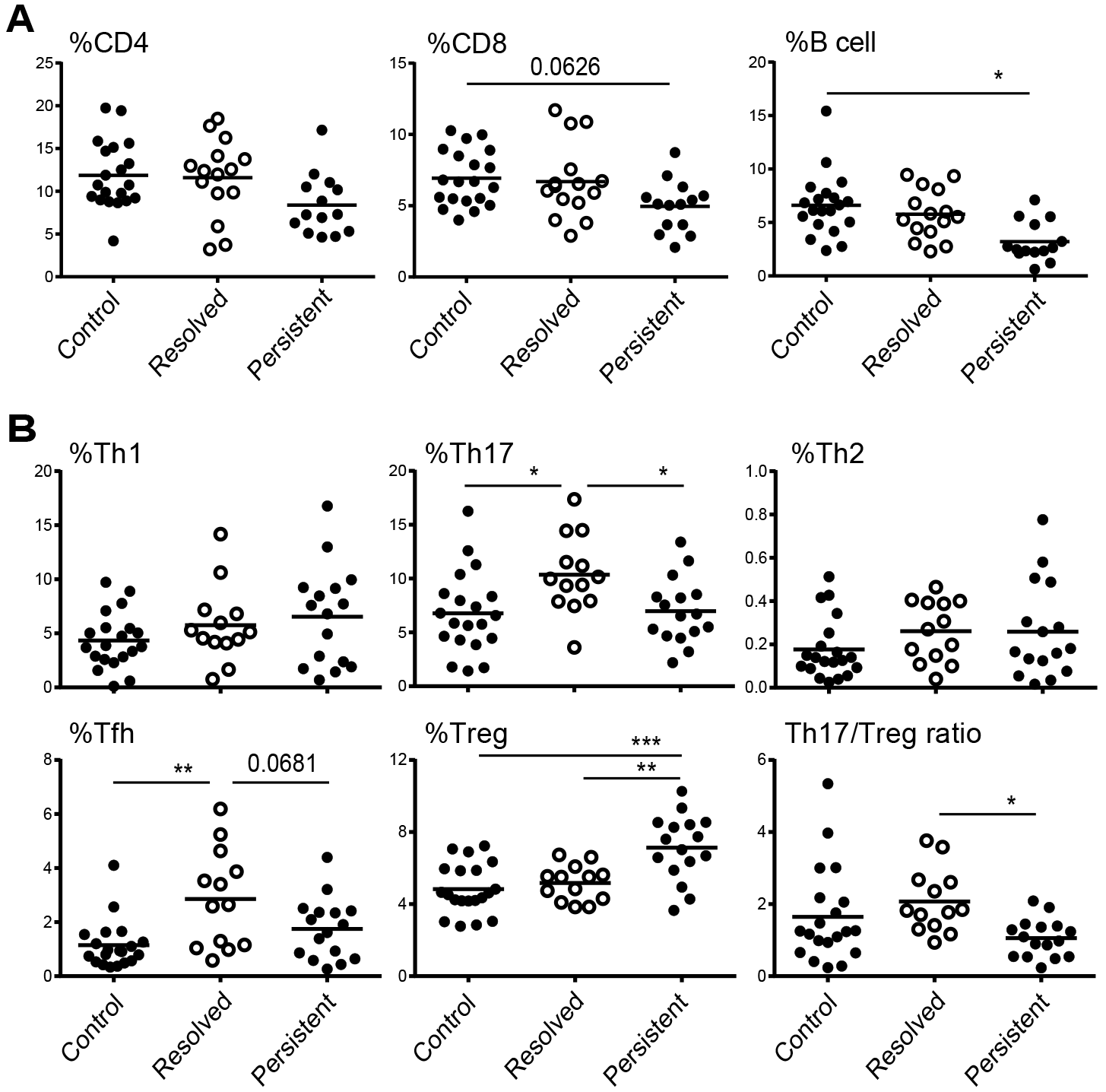
T helper cells determine disease outcome. PBMCs from acute disease were incubated with antibodies and the percentage of adaptive immune populations (A) and T helpers (B) determined by flow cytometry. T helper subset frequencies are represented as percentage positive within the CD4^+^CD45^+^ gate. Samples were separated by healthy, resolved or persistent infection. Long lines indicate the mean, and each dot represents an individual patient. ANOVA with Bonferroni correction was used to compare multiple groups with a 95% confidence interval; * p<0.05, ** p<0.005, *** p<0.0005.

Immune cells secrete inflammatory and effector cytokines to induce cell migration, differentiation and function. We evaluated the concentration of 26 inflammatory and Th cytokines (Figure 3A). As cytokines were evaluated in serum, and not from stimulated cell supernatants, most cytokines were expressed at just above the limit of detection (LOD). IL-1α, IL-1β, IL-5, IL-12p70, IL-21 and IL-22 were below the LOD for all patients (not shown). Significant differences in IL-6, IL-18 and IL-12, and mild difference in IL-10, were observed between patients with resolved and persistent disease, and all of these were increased relative to healthy controls (Figure 3B). IL-6, IL-18 and IL-12 are produced by antigen presenting cells and promote T effector differentiation. IL-10, an immunosuppressive cytokine, is expressed by several immune populations, including Tregs. IL-18 inhibits Th17 and promotes Treg differentiation and function [25, 26].

**Figure 3.**
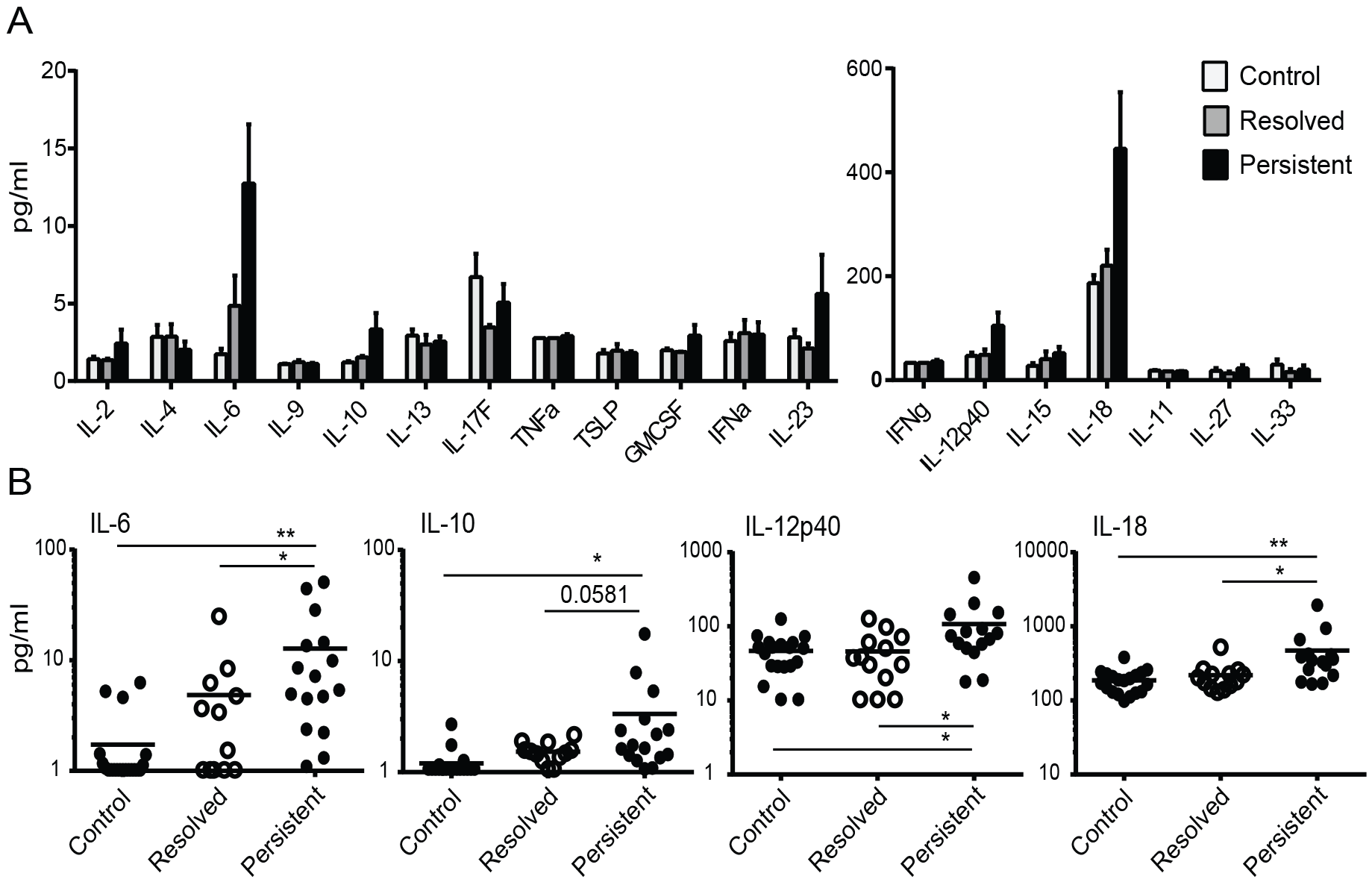
Serum cytokine data. Serum was incubated with a panel of cytokine antibodies and the concentration determined by flow cytometric analysis. Samples were separated by healthy, resolved or persistent infection. A) Cytokines are shown with the mean and SEM. IL-1α, IL-1β, IL-5, IL-12p70, IL-17A, IL-21, and IL-22 were below the LOD, and are not shown. B) Individual cytokines are shown with each dot representing an individual patient. Samples at or below the LOD are shown at the LOD. ANOVA, then a Fisher’s Least Significant Difference was performed; * p<0.05, ** p<0.005.

### Predictive biomarkers of disease outcome identified

To identify predictive biomarkers we performed principle component analysis (PCA) using all non-diagnostic clinical, immune population and cytokine parameters. PCA analysis separated patients from healthy controls, and inpatients from outpatients, but was unable to distinguish patients with resolved versus persistent disease (Figure 4A). PCA comparison, using all parameters, only described 31.9% of variation in the hospital status data and 26.3% of variation in the disease outcome data, indicating that some parameters are too variable to include as a means to distinguish patients. Chi-square automatic interaction detection (CHAID), a decision tree technique, was used to define a predictive method for disease outcome. Treg frequency had the greatest impact on disease outcome of all clinical and immune parameters (Figure 4B). Treg frequency at diagnosis distinguished disease outcome in 79.3% of patients (11/13 resolved and 12/16 persistent disease). Eosinophil, neutrophil and Th17 numbers further defined patients within these disease outcomes (not shown). PCA using the parameters identified by CHAID in the first two levels of separation (%CD4^+^Treg, eosinophils and neutrophils), predicted disease outcome in 89.7% (26/29) of patients.

**Figure 4.**
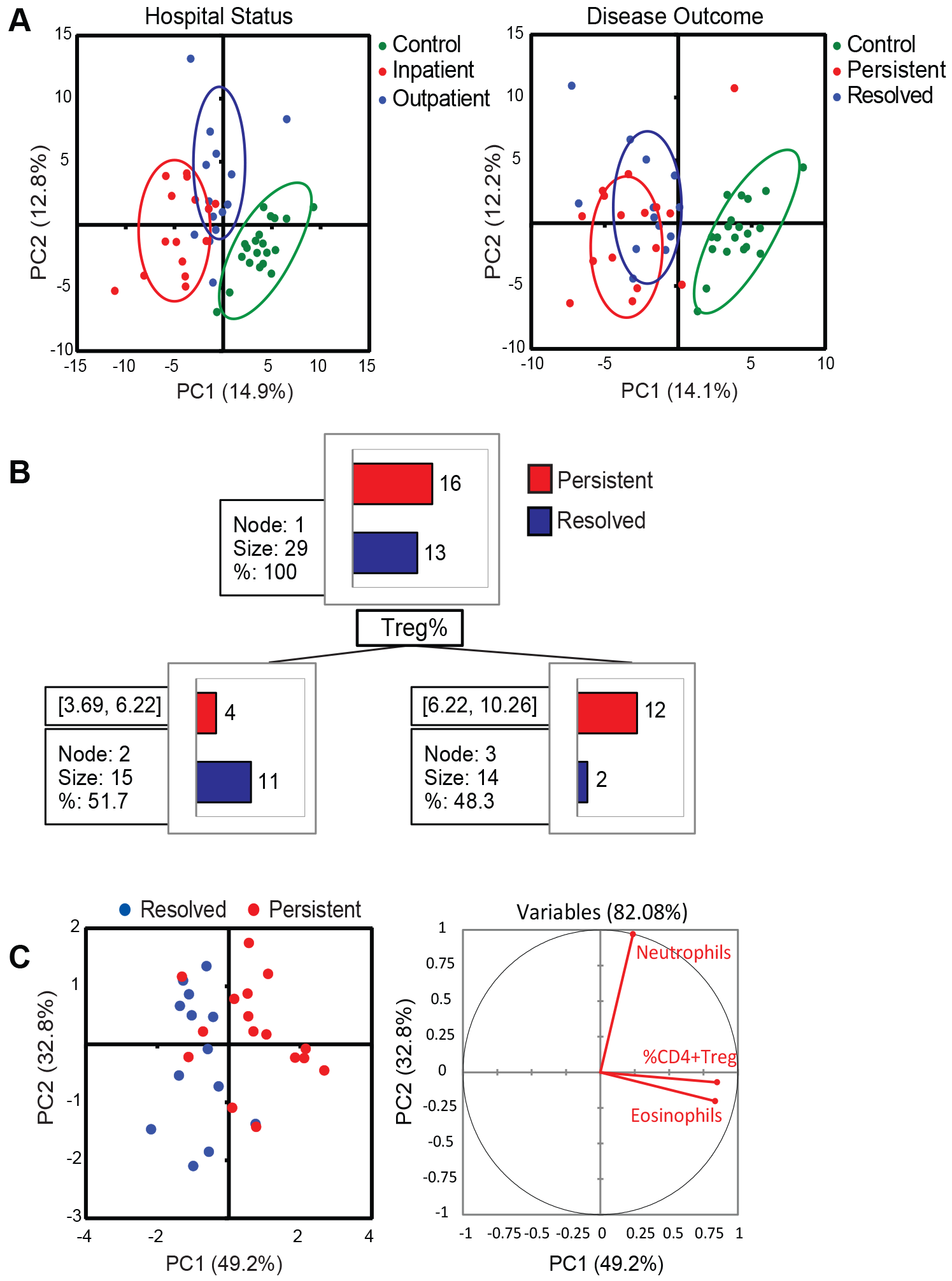
PCA and CHAID analysis reveals a relationship between Treg frequency and *Coccidioides* infection. A) PCA using 119 clinical and flow cytometry parameters of controls, inpatients and outpatients (left graph), or controls, resolved and persistent disease (right graph). B) CHAID analysis on persistent and resolved patients identifies a relationship between %CD4^+^ Treg and disease outcome. C) PCA analysis on persistent and resolved patients using eosinophil and neutrophil numbers, and %CD4^+^ Treg parameters.

### Unbiased high-dimensional flow cytometry analysis reveals CCR5 as a biomarker of persistent disease

We applied high-dimensional flow cytometry analysis tools to identify cellular populations that predict disease outcome. We utilized CITRUS, a tool that identifies stratifying cellular signatures that differ between grouped data [27]. After pre-gating CD4^+^CD45^+^ T cells, we utilized CD127 (IL-7Rα), CXCR5 (CD185), CCR3 (CD193), CCR5 (CD195), CCR6 (CD196), CD25 (IL-2Rα), and HLA-DR to cluster the data. Comparing resolved versus persistent disease patients, CITRUS identified several cell populations that stratified the data at a 1% false discovery rate, including one population representing Tregs (Figure 5A). CITRUS clusters revealed CCR5 (CD195) as expressed more highly in persistent patients; specifically, elevated CCR5 stratified Tregs. The frequency of the CCR5+Treg population was higher in persistent than resolved disease (Figure 5B). CCR5 is critical for Treg migration into tissues [28].

**Figure 5.**
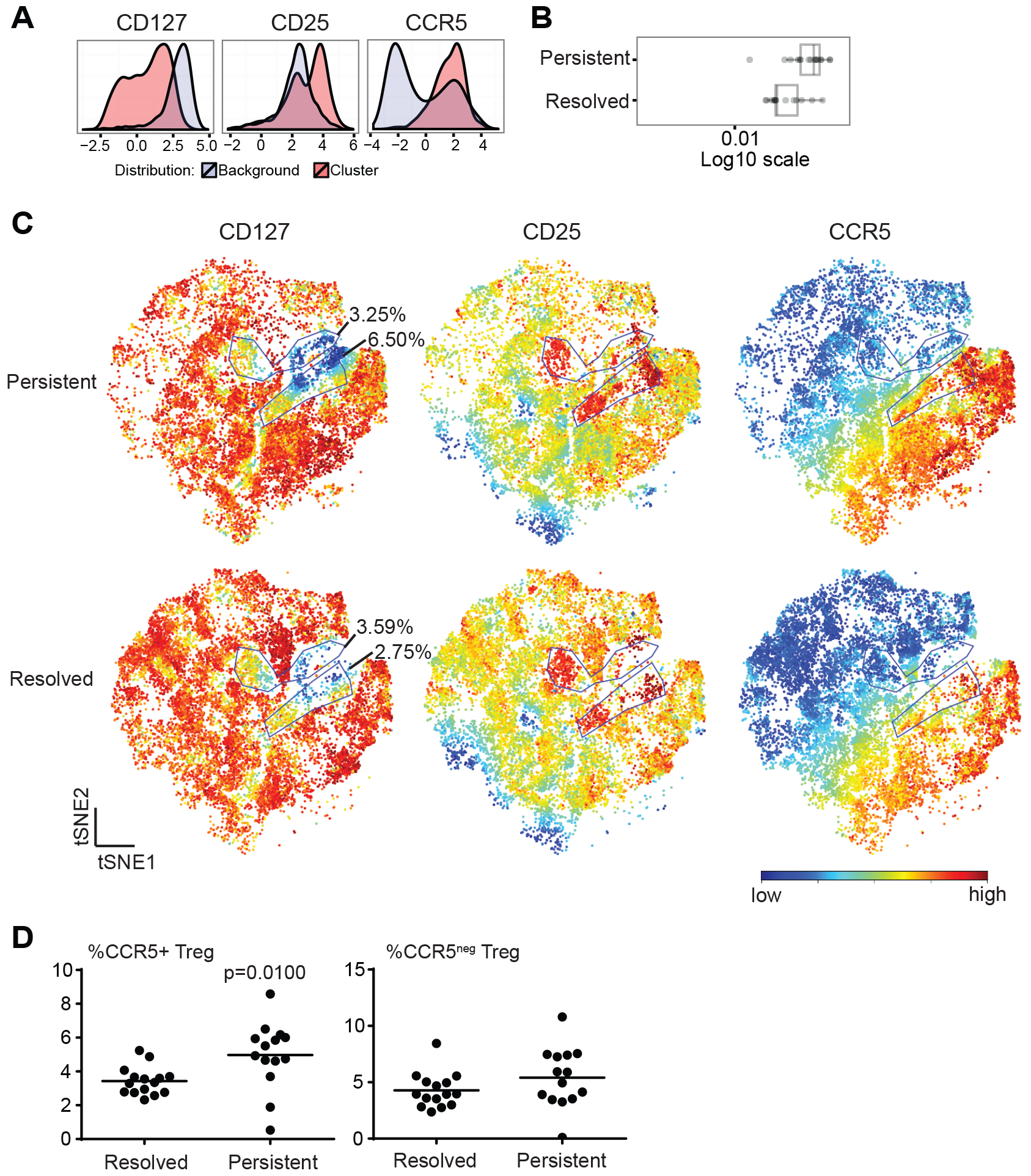
CCR5^+^ Tregs are a significant predictor of disease outcome. CITRUS analysis was performed comparing persistent and resolved patients. Within the CD4^+^CD45^+^ population, CITRUS identified several differential cell clusters that separated patient groups. CD127, CCR5, and CD25 expression is shown for the Treg cluster (A). Box plots display the frequencies of the significant Treg cluster between persistent versus resolved patients demonstrating this cluster is elevated in patients who go on to have persistent infection (B). (C) viSNE heatmaps were generated for CD25, CD127, and CCR5 on two representative patients. Gates were drawn around two Treg (CD4^+^CD45^+^CD127^low^CD25^+^) clusters based on CCR5 expression. %CCR5^+^ Tregs and %CCR5^neg^ Tregs was determined across all samples (D). An unpaired student’s t-test was performed.

Performing viSNE, a dimensionality reduction and data visualization tool [29], we confirmed a region of Tregs that could be divided on CCR5 expression. (Figure 5C) [29, 30]. CCR5^+^Tregs were significantly elevated in patients with persistent disease as compared to patients who resolved infection, while CCR5^neg^Treg frequency was unchanged (Figure 5D). High-dimensional analysis confirms a Treg disparity based on disease outcome, and reveals CCR5 as a functional marker associated with chronicity.

## DISCUSSION

We identified several clinical measurements and immune populations that together provide potential biomarkers for predicting disease outcome in *Coccidioides*-infected patients. Elevated Treg frequency was the major indicator of persistent infection. Most patients with persistent disease remain on anti-fungal treatments for months to years, with risk of fungal reactivation upon therapy discontinuation. Understanding host immune responses leading to dissemination, persistent infection or resolution of infection may allow clinicians to identify patients with persistent infection earlier.

Patients with persistent disease appear to develop an inappropriately prolonged innate immune response, and ineffective adaptive immune activation to *Coccidioides* infection. Expanded neutrophils and eosinophils indicate that the innate immune system in these patients mounts an inflammatory response, in contrast to reduced lymphocyte frequencies. As lymphocytes and appropriate CD4^+^ T cell responses are required for immune control of *Coccidioides*, patients with persistent infection likely do not have sufficient activated lymphocyte numbers to control infection. Resolution of infection appears to require a Th17, but not a Th1, response that is reduced in patients with persistent infection.

The four patients who developed disseminated infection had significantly reduced alanine transaminase (ALT) levels compared to resolved patients (p=0.0014; not shown), indicating liver dysfunction not previously described in disseminated coccidioidomycosis, although elevated aspartate aminotransferase:ALT ratio is observed in disseminated histoplasmosis [31]. We observed trends indicating reduced adaptive immune responses (reduced CD4^+^, CD8^+^, B, and Th17 cells) and ineffective innate immune responses (elevated classical monocytes) [32]. Further study of disseminated infection is needed to clarify the immune responses in disseminated coccidioidomycosis.

Tregs suppress T cell activation and effector mechanisms [30]. Enhanced Treg ratios and suppressive function may block appropriate adaptive immune responses, enabling prolonged fungal infection, resulting in persistent infection. Elevated Treg frequency provided the best biomarker for identifying these patients at diagnosis, suggesting that Tregs may suppress appropriate immune responses during *Coccidioides* infection. Alternatively, as Tregs and Th17 cells differentiate through reciprocal pathways, Tregs may be generated at the expense of Th17 cells required for fungal clearance, resulting in persistent infection [15]. These patients may have naturally high Treg frequencies before infection, or as suggested by the significant elevation in Treg frequency relative to healthy controls, may ineffectively induce peripheral Tregs instead of Th17 effectors. Serum cytokine levels in patients with persistent infection further support the possibility of inappropriate T cell differentiation. IL-18 inhibits Th17 and enhances Treg differentiation and function, which could explain persistent infection in these patients [25, 26].

In *Paracocidioides brasiliensis* infection Treg frequency and suppressive function is enhanced relative to controls [18]. Our results support a similar finding in *Coccidioides* infection, with higher Treg frequencies indicative of persistent infection, and markers of suppressive Treg function. CITRUS analysis revealed a higher CCR5^+^Treg frequency in persistent infection. CCR5^+^Tregs have enhanced suppressive capacity compared to CCR5^neg^Tregs [33]. CCR5-deficient mice have an enhanced ability to combat *Paracoccidioides brasiliensis* infection, and elevated Th17 response to *Histoplasma* infection, resulting in a Treg/Th17 imbalance [34, 35]. We speculate that Treg expansion and elevated functionality prevent appropriate Th17 effector responses allowing persistent *Coccidioides* infection.

The elevated Treg frequency and potential suppressive functionality is consistent with our understanding of Tregs in infection, and with the inverse relationship in Th17 and Treg differentiation. It is tempting to propose that manipulating Tregs or Th17 cells could improve prognosis for *Coccidioides*-infected patients. Further predictive modeling is needed to confirm our findings, and better understand the Treg phenotype during infection. To our knowledge, this is the first pediatric study demonstrating higher Treg and lower Th17 levels in patients who cannot clear infection. These markers could identify patients earlier in their disease course who would benefit from prolonged treatment course, and close monitoring for disease relapse once off therapy.

## FUNDING

This work was funded by the Blum Center; Valley Children’s Healthcare; the University of California Merced Health Sciences Research Institute; and by a gift from Dr. E.W. & Dorothy Bizzini.

## ACKNOWLEDGMENTS

We thank Padma Desai and Christine Banda for clinical enrollment support, our patients for participating in the study, Dr. Larry Fong for helpful discussions, the UC Merced Stem Cell Instrumentation Foundry for their assistance in flow cytometry design, and the UC Merced Biostatistics and Data Support Core for consultation on statistical methods.

## Notes

*Conflict of interest*. The authors have nothing to disclose.

*Financial support*.This work was funded by the Blum Center; Valley Children’s Healthcare; the University of California Merced Health Sciences Research Institute; and by a gift from Dr. E.W. & Dorothy Bizzini.

*Presentations*. This work has been presented at the 7th International Coccidioidomycosis Symposium and at the 2018 Federation of Clinical Immunology Societies Annual Meeting.

